# Fine-grained, Nonlinear Image Registration of Live Cell Movies Reveals Spatiotemporal Organization of Diffuse Molecular Processes

**DOI:** 10.1101/2021.11.22.469497

**Authors:** Xuexia Jiang, Tadamoto Isogai, Joseph Chi, Gaudenz Danuser

## Abstract

We present an application of non-linear Image registration that allows spatiotemporal analysis of extremely noisy and diffuse molecular processes across the entire cell. To produce meaningful local tracking of the spatially coherent portion of diffuse protein dynamics, we improved upon existing non-linear image registration to compensate for cell movement and deformation. The registration relies on a subcellular fiducial marker, a cell motion mask, and a topological regularization that enforces diffeomorphism on the registration without significant loss of granularity. We demonstrate the potential of this approach in conjunction with stochastic time-series analysis through the discovery of distinct zones of coherent Profillin dynamics in symmetry-breaking U2OS cells. Further analysis of the resulting Profilin dynamics revealed strong relationships with the underlying actin organization. This study thus provides a framework for extracting functional interactions between cell morphodynamics, protein distributions, and signaling in cells undergoing continuous shape changes.

**Author Summary:** By adapting optical flow based nonlinear image registration we created a method specific for live cell movies that preserves the dynamics of a signal of interest by remapping using a separate measurable subcellular location fiducial. This is an extension as well on our lab’s previously published method of cell edge-based referencing of subcellular locations that was incapable of extracting interpretable subcellular time series more than a few microns away from the cell edge. We showed that our method overcomes this key limitation and allows sampling of subcellular time series from every subcellular location through our discovery of organized profilin dynamics in moving cell and that these profilin dynamics are related to actin dynamics due to their ability to bind growing actin structures likely through actin polymerizing factors. Most importantly, our method is applicable to discovering subcellular organization and coordination in a widely used form of live cell microscopy data that hitherto has been largely limited to anecdotal analysis.

## Introduction

Time-series analysis of live cell movies can quantify functional interactions between proteins in the context of complex regulatory networks [1–4]. Utilization of these statistical tools requires extraction of time-series in a cell-centric frame of reference that retains the locations of molecular interactions over time despite cellular movements and shape variation. For cell biology this means following these locations through large morphological deformations such as protrusions, retractions, organelle translocations, and cytoskeletal rearrangements. Because of the difficulty in tracking subcellular locations, most analyses of live cell movies have remained incomplete and superficial, limiting the ability to test quantitative models of cell behavior.

Historical solutions to the extraction of temporal information in one location can be broken down into three categories: manual sampling (i.e. kymographs) [5, 6], cell edge propagated sampling [1], and experimental approaches that limit cell deformation [7]. Kymographs, while intuitive and easily used, are by nature incapable of simultaneously handling the entire cell and even locally tend to introduce significant artifacts to the time series as they do generally not follow cell deformation [5, 6]. The introduction of edge propagated sampling has overcome some of these issues, in principle [1, 8]. However, while registration of locations near the cell edge reveal expected interactions between proteins [1, 2, 9], time series at locations deeper inside the cell reveal a convolution of the real molecular dynamics with cell motion, leading to uninterpretable data typically a mere few microns away from the cell edge. Experimental constraints such as micropatterning try to overcome this problem by fixing the cell footprint to a particular shape [10], but they do not necessarily limit the subcellular reorganization of molecular activities. More importantly these constraints often introduce harsh perturbations to the cell architecture, which obscure many of the natural cellular behaviors. A more promising approach is the use of image processing to constrain the footprint of the cell across time through nonlinear registration. This approach can either utilize principles of optimal transport or optical flow.

Optimal transport methods find the minimum movement necessary to redistribute the intensity from an input image to the intensity of a target image [11–13]. This is well suited to registration of images collected from multiple samples since minimal displacement is the most appropriate assumption without real displacements between the two images to approximate. Prior work at the cellular scale utilizing optimal transport principles to track time series in live neutrophils focused on a stereotypical set of behaviors, i.e. the formation of an immunological synapse attacking a model bead [11]. Neutrophils, whose morphology differed too much from the stereotype, were dismissed. Although in this study the authors compiled a great number of signaling time-series from morphologically diverse neutrophils, the analysis was focused on the narrow synapse region with almost rigid geometry. Moreover, it was limited to low spatial resolution, which softened the requirement to align time series with detailed variations of the synapse shape.

Optical flow-based methods also work under the assumption of intensity conservation but estimate local displacements based on local intensity differences and the observed intensity gradient [14–16]. These methods have been applied to great success in registration of functional brain studies that also seek to extract time-series with high spatial granularity [17]. However, because of the aperture problem and image noise, Optical Flow computation is ill-posed and thus requires domain-specific regularization techniques. This has previously hindered the application of these methods to live cell imaging, which presents a much harder registration problem than brain imaging. Live cell time-lapse sequences are often characterized by large geometric deformations and image structures that are captured at the limit of resolution. However, proper constraints in principle allow approximation of true underlying motion [14]. Here we present an optical flow-based approach with a regularization scheme tailored for cell biological applications. The method enables image registration of cells throughout extended live movies. We then illustrate how the result enables the extraction of local time series across the entire footprint of arbitrarily deforming cells. To demonstrate the unprecedented potential of this time series extraction pipeline we chose to analyze the dynamics of profilin, which is a diffuse cytoplasmic protein and a component of the actin polymerization machinery, whose interactions with actin cannot be interpreted without registration of the live cell movies.

## Results

### Subcellular protein location referencing reveals spatiotemporal dynamics in live cell movies

To extract time-series from live cell movies all frames must be mapped from the lab-centric frame of reference of the image detector into a cell-centric frame of reference, where any subcellular location covers a relevant time-series. This is an ill-posed problem since we cannot directly observe all physical and chemical processes that govern the transport of a labeled protein (signal of interest) over time, including the changes in cell morphology itself. Nevertheless, by using an approximation of these processes using optical flow principles we can appreciate subcellular dynamics approaching the imaging resolution.

Among the various techniques for computing optical flow, Thirion’s demons has gained popularity because of fast computation speed, intuitive approach, and ease of adjustment of the regularization to diverse data sets [18–20]. Thirion’s demons define correspondence between two images following locally the intensity gradient to approximate the “diffusion” process that took place between the two images [18]. The procedure thus makes minimal assumptions about the data and performs well when a movie is sampled quickly relative to the changes in morphology, a shared prerequisite for the intended goal of time-series analysis.

Our goal with this presented pipeline is to facilitate the high-content analysis of subcellular patterns of dynamic molecular activities visualized via fluorescently labeled proteins in 2D movies of single cells. We conceptualize the cell as a dense space where the location of a molecular activity or concentration, referred to as the signal of interest, is coupled to the morphodynamics of the cell (Fig. 1a, second row). The morphodynamics is observed by a separate probe, referred to as the location fiducial (Fig. 1a, top row). Using the location fiducial, we apply nonlinear image registration techniques with the objective of matching all location fiducial images in the movie as closely as possible to a reference frame (Movie 1). The reference frame can be any image of the same location fiducial, including an image of another cell. Without losing the power of generalizing the pipeline, we will focus here on demonstrating the scenario where the middle frame of a movie is used as the reference frame. We then remap images of the corresponding signal of interest using the deformations that produced closely matched fiducial images to compile a movie of the signal of interest with a fixed cell shape (Fig. 1a, third and fourth row). The resulting movie exhibits now in a fixed cell shape dynamics of the signal of interest that were not coupled to changes in the location fiducial (Movie 2).

**Figure 1.**
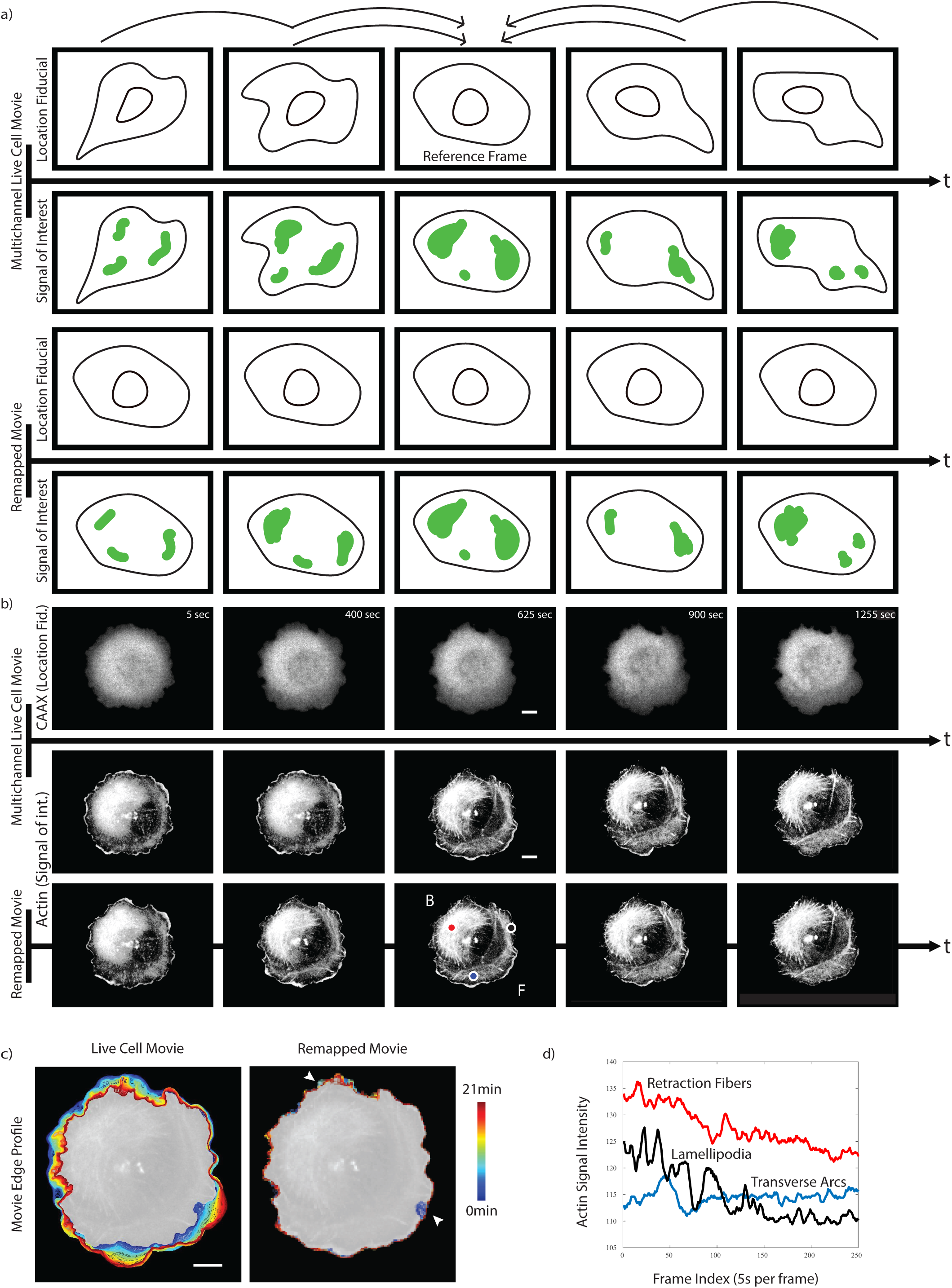
Registration of a signal of interest captured by a live cell time-lapse sequence using a co-imaged location fiducial. a) Illustration of the pipeline. Top row: The pipeline establishes a frame-by-frame diffeomorphism between all images of a location fiducial time lapse sequence and a reference frame. Without loss of generality, all analyses in this paper rely on the center frame of the sequence as the reference frame. Bottom row: The diffeomorphic functions are applied to remapping a signal of interest from the original time lapse sequence to a new time lapse sequence with a rigid geometry defined by the location fiducial reference. The dynamics of the signal of interest is conserved by the remapping process. b) Application of the pipeline to a cell undergoing symmetry breaking, i.e. it transforms from a round geometry to a geometry with front (F)-back (B) polarity. The data includes a Halo-CAAX and mNG-actin time lapse sequence as a location fiducial and as the signal of interest, respectively. Colored points in the center frame of the remapped actin signal indicate the position of three sampling positions illustrated in d. c) Cell edge color-coded from blue (early time points) to red (late time points) in the geometry of the original time lapse sequence and in the geometry of the reference frame. With the exception of two regions with temporarily strong ruffling activity (arrowheads), the remapping produces a stationary cell edge. d) The mapping into a rigid frame of reference permits straightforward sampling of time series of the signal of interest. The traces indicate actin signal fluctuations at stereotypical sites of interest: Red – retraction fibers, Blue – transverse arcs, Black – lamellipodia. All scale bars 10 μm.

The implemented pipeline rests on important assumptions about the data and signal of interest: i) There should be fast sampling of the movie such that the sampling rate is below the Nyquist limit of the biological behavior of interest. This also means that we expect small shape changes between timepoints but large shape changes can occur over the course of the movie. ii) The cell should sit flat in 2D and should not move out of view for the duration of the analysis. While the cell can contact other cells, they should not move below or above other cells as in 2D this produces a region of high intensity in the fiducial channel that will be separated by the remapping process. iii) Most importantly we assume that subcellular motion is faithfully represented by a location fiducial marker.

Fig. 1b introduces the pipeline on the example of a polarizing U2OS cell. Over the course of approx. 20 minutes the cell changes from a rounded shape to a canonical migratory shape with a leading edge. We use Actin as the signal of interest and a CAAX membrane marker as the location fiducial. The method eliminates shape deformation except for spurious artifacts in two peripheral regions of erratic ruffle formation where cell edge tracking fails (Fig. 1c, Movie 3).

After remapping, we can readily extract time-series of the signal of interest at any subcellular location (Fig. 1d, positions of lamellipodia, retraction fiber, and transverse arcs are indicated in the reference time point of Fig. 1b 3^rd^ row). Of note, in this example the actin intensity is the signal of interest, and the remapping over time is accomplished based on a deformation field derived from a diffuse signal. Nonetheless, fine fibrous structure both in the front and in the back of the cell are preserved (Fig. 1b 3^rd^ row), indicating a precise estimation of the deformation field over time. Visually we can see for a stereotypical cell waviness in actin filaments indicative of the fact that motion of these large filaments is not identical to motion of a diffuse membrane marker. This breakdown in our assumptions and any imperfections in registration precludes extraction of dynamics at the imaging resolution (120 nm). The rest of this paper will attempt to quantitatively establish that our method produces time-series relevant for micron scale behaviors.

### Algorithm basis for fitting deformation fields

The Thirion’s demons algorithm framed image registration as a diffusion process whereby local forces, inspired by optical flow equations, pushed a moving image onto a target according to local characteristics of the images [18, 19]. The algorithm takes as input a moving image (movie frames) and a target image (the reference frame) and outputs a deformation that remaps the moving image through interpolation to be as close to identical to the target image as possible. The algorithm alternates between calculating the forces based on intensity differences and spatially regularizing the forces with a Gaussian kernel. For a given coordinate, let **m** be the local intensity of the moving image and **f** be the local intensity of the target image. The local displacement **u** is given by eq1.

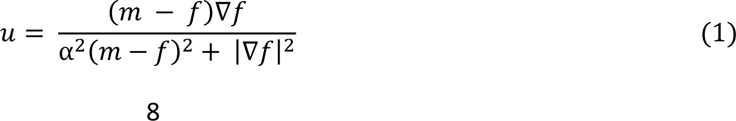

Here, **∇f** denotes the gradient of the target image intensity and **α** is an optional throttling term usually set to limit the maximal displacement calculated at each iteration to 1 pixel. The matrix of these calculated displacements, **U**, is then smoothed by a Gaussian kernel **K_diff_** and the entire process repeated **n** times until a registered image is achieved.

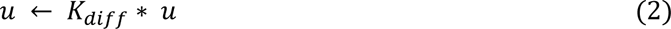

The solution for any given two images depends on the setting of **α** and **K_diff,_** which determines the influence of small, local structures on the image matching. Since our goal was to approximate observable motion in cells, we made a series of cell biology specific adaptations to Therion’s demons described next.

To determine the **n** iterations we picked a high motion movie in our dataset and determined the cell mask deviation over the iterations (Fig. 2c). The iteration number **n = 200** was selected such that we get near pixel alignment. From prior knowledge we expect the observable motion at the cell edge to be far greater than the motion in the cell interior [21]. This value is fixed in the current software distribution, but can be readily adjusted by a user.

**Figure 2.**
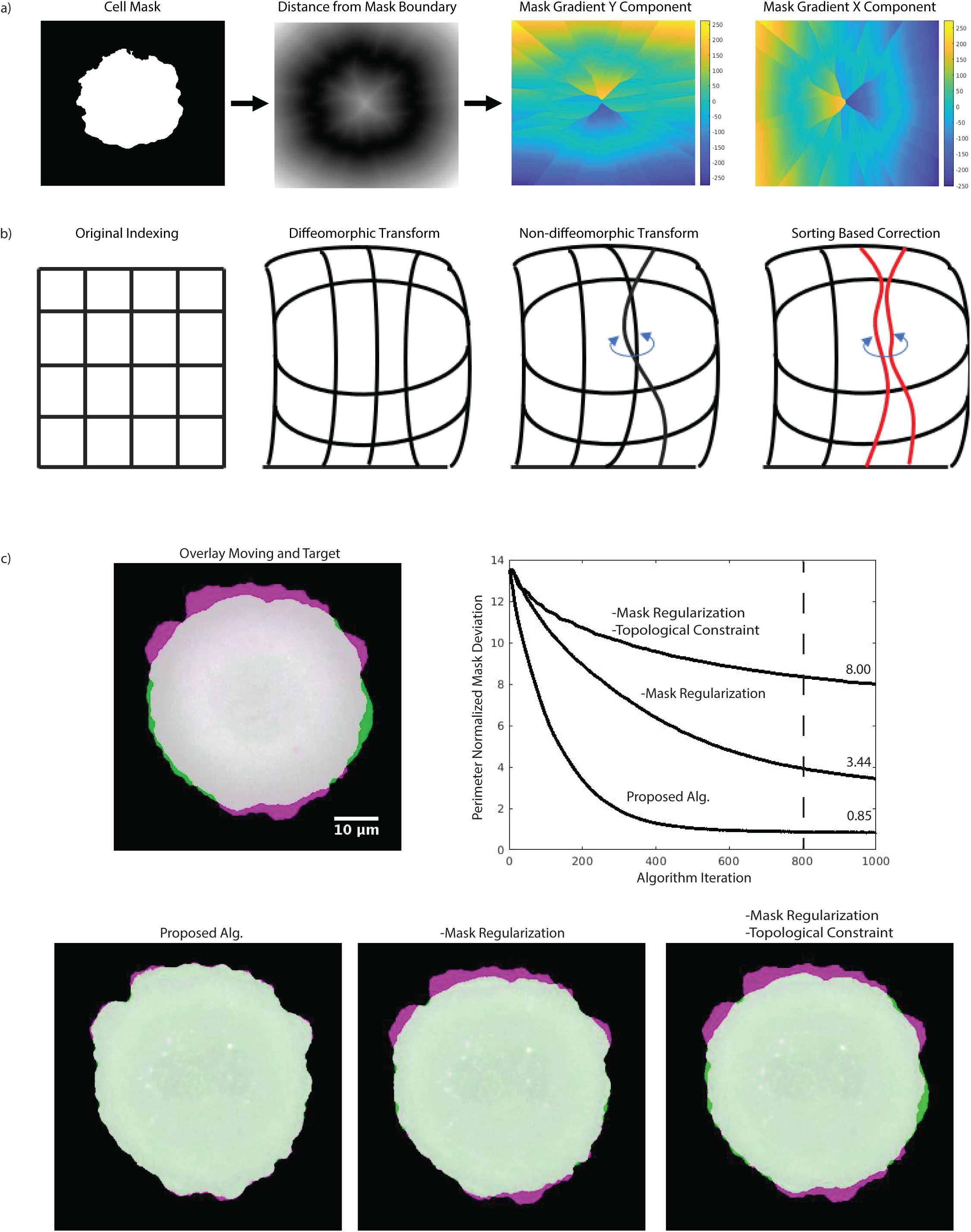
Computational elements for the remapping process. a) Extraction of mask gradient (∇fmask). Based on a cell mask, we calculate for each pixel the distance to the nearest cell boundary element for both the cell-interior and cell-exterior spaces. The image gradient for this distance transform defines ∇fmask. b) Diffeomorphism constraint. From left to right, illustrations of interpolation fields, where edges indicate the sampling position of an input image to remap onto a target. The mesh diagram shows displacements as deviation from a square mesh (1^st^ diagram from left). A diffeomorphic transform (2^nd^ diagram) is a smooth manifold and a break in diffeomorphism (3^rd^ diagram) is seen as a crossing of mesh edges. The mesh vertices are coordinates for resampling an image. By sorting these coordinates in sequential order, we correct for a break in diffeomorphism (4^th^ diagram). c) Effect of algorithm components on cell edge alignment. Accuracy of registration over n iterations is indicated by the area of mismatch (green and purple) between moving cell and target, normalized by the target cell perimeter. Removing the mask regularization and topological constraint reduces both the rate of convergence and the final accuracy. The dashed line indicates the iteration stop point for a visualization of the registration results (bottom row). The proposed algorithm gives a near pixel perfect registration of the two images. Removing mask regularization largely reduces the rate of convergence. Removing both mask regularization and topological constraint causes an inability to capture small protrusions likely due to image noise.

To remap a sequence of frames ending at the final reference frame, we change the target image after **n** iterations to the next frame in the sequence until we fit the deformation to the reference frame after **n_tot_** iterations (Fig. 1a) so that farther time points with more frames in sequence receive more passes through the algorithm.

Biological data contains much higher noise and lower dynamic range than photography images for which the original framework has been developed. In the extreme case the cell exhibits a checkerboard like pattern (e.g. peripheral branched actin in Fig. 1b) and background outside the cell that is close to the intensity inside the cell. This creates a scenario where the result is highly dependent on **α** and **K_diff_** values where low values will cause the algorithm to get stuck in local minima and large values will cause the algorithm to not register small protrusions with known biological importance. To solve this, we introduced a mask regularization component where the mask is a segmentation of the cell separating the cell from background and other cells in the field of view:

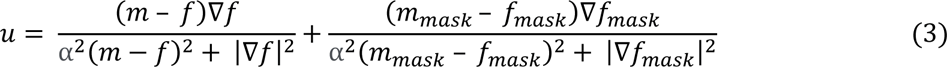

Here **m_mask_** is the location of the segmentation mask in the moving image and **f_mask_** is the location of the mask in the target image. We derive the mask gradient **∇f_mask_** over the entire cell footprint from the gradient of the map of shortest distances to the cell edge (Fig 2a). In practice the mask regularization component is non-zero where the two image segmentations do not overlap. Thus, it guides the registration process to an overlay of the two cell images moving the registration out of local minima. In presence of a high-quality segmentation of the cell outline, the regularization component is independent of the noise of the fiduciary signal throughout the cell. This permits us to set **K_diff_** based on the expected diffusion distance between time steps as opposed to an arbitrary number for better registrations. We then set **α** to limit displacements to 1 pixel, but higher limits can be used to quickly test suitability of input data for this pipeline (not shown).

The original publication of Thirion’s demons presented a way to enforce diffeomorphism based in Lie group theory.[18] The approach broke each displacement into a series of smaller steps based on the magnitude of the displacement and smoothed for each of these steps. This approach tends to be trapped in local minima, especially with a parameter selection meant to capture the granularity in cell data [20]. Instead, we chose to use a topological regularization through sorting. Given the 2D matrix of unsorted displacements **U**, we compute **U_sorted_** as a matrix such that:

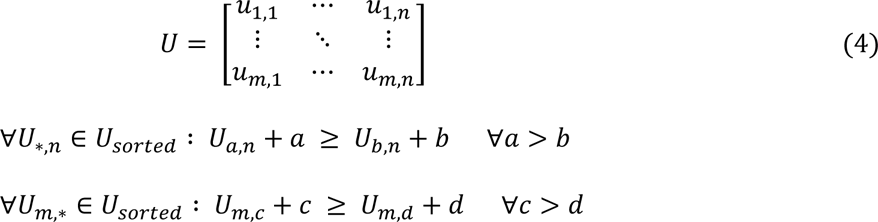

To illustrate how this process works, a deformation field **U** is transformed into a matrix of indices for interpolation as per the image registration process. This can be represented in mesh format (Fig. 2b, panels i and ii) where square grids indicate an identity transform of the original image and non-square grids indicate either local dilation or contraction. A break in diffeomorphism occurs when indices are out of order leading to a crossing or fold in the mesh representation (Fig. 2b, panel iii). Per iteration the sorting procedure corrects violations of the diffeomorphism (Fig. 2b, panel iv) by reordering the indices in the matrix. In practice, diffeomorphism breaks occur when either a difficult checkerboard pattern or noise continue to push the optimization towards a registration with crossings. These regions through our topological sorting will converge up to the step limit imposed by **α** and **K_diff_.**

To illustrate the power of our approach we chose a U2OS cell undergoing a stereotypical isotropic spreading process and remapped 2 distant time points while setting **K_diff_** to a non-diffusion related low value of 0.5. This condition deviates from our expectation of fast sampled changes but allows easy interpretation of the impact of our modification to the original Thirion’s demons and shows that our approach can handle more extreme morphology changes, such as those found in long time-lapse imaging of cells. In combination, the motion mask and sorting based regularization permit near-pixel perfect edge alignment between two distant time points (Fig. 2c). Without these constraints, a typical remapping can exhibit perimeter-normalized boundary deviations (total pixel difference in image masks/length of perimeter in pixels) larger than 8 pixels. This can correspond to losing an entire protrusion event. Our adapted algorithm additionally requires fewer iterations to converge to an accurate edge alignment, showing that computational efficiency is not traded for accuracy.

### Evaluation of Algorithm by Image Signal Remapping

Our approach relies on a subcellular location fiducial. Generally, this requires an additional fluorescence channel for live cell imaging besides the signal of interest. The use of a dedicated channel only for image alignment can be quite cumbersome and increase photo-toxicity. We therefore tested the use of a down-sampled signal of interest as a location fiducial. Since the goal of the image alignment is the extraction of informative time-series, we can treat the signal of interest as a mixture of high spatial frequency signals describing local molecular activity and low spatial frequency signals describing the dynamics of subcellular molecular organization at a coarser scale. Assuming separability of spatial frequencies and a known frequency cutoff, we can use the low frequency bands (down sampled images) for registration without artificially flattening the informative high frequency signals. In principle the combination of fast sampling, diffeomorphism constraint, and mask regularization limits the space of possible remapping differences. Nevertheless, the choice of location fiducial impacts the remapping results and we sought to quantify the impact of location fiducial choice.

Real world photography and medical image registration use landmark comparison and reconstruction of computationally distorted images to measure performance. Neither is a feasible metric for our live cell movies as we have no ground truth except for specific anecdotal examples. Moreover, a simulated distortion can arbitrarily favor a particular fiducial choice. We therefore introduce the ‘to-target transform’ and the ‘half-distance transform’ accuracies as alternative performance measures for the remapping quality. For both measures, we first remapped the signal of interest from moving image to the target image using location fiducial deformation fields over two frames. For the to-target transform accuracy we then asked how similar the remapped signal of interest is in comparison to the imaged signal of interest in the target image. The proximity between remapped and target signals of interest was determined by the average pixelwise squared intensity difference. This metric quantifies how much of the dynamics is captured by the location fiducial and our remapping process. For the half-distance transform accuracy, we divided the magnitude of the deformation field over two frames in half and compared the thus remapped signal of interest to the signal of interest in the skipped frame (Fig. 3c). This metric quantifies how well the approximations of a diffusion process underlying the remapping capture the real dynamics.

**Figure 3.**
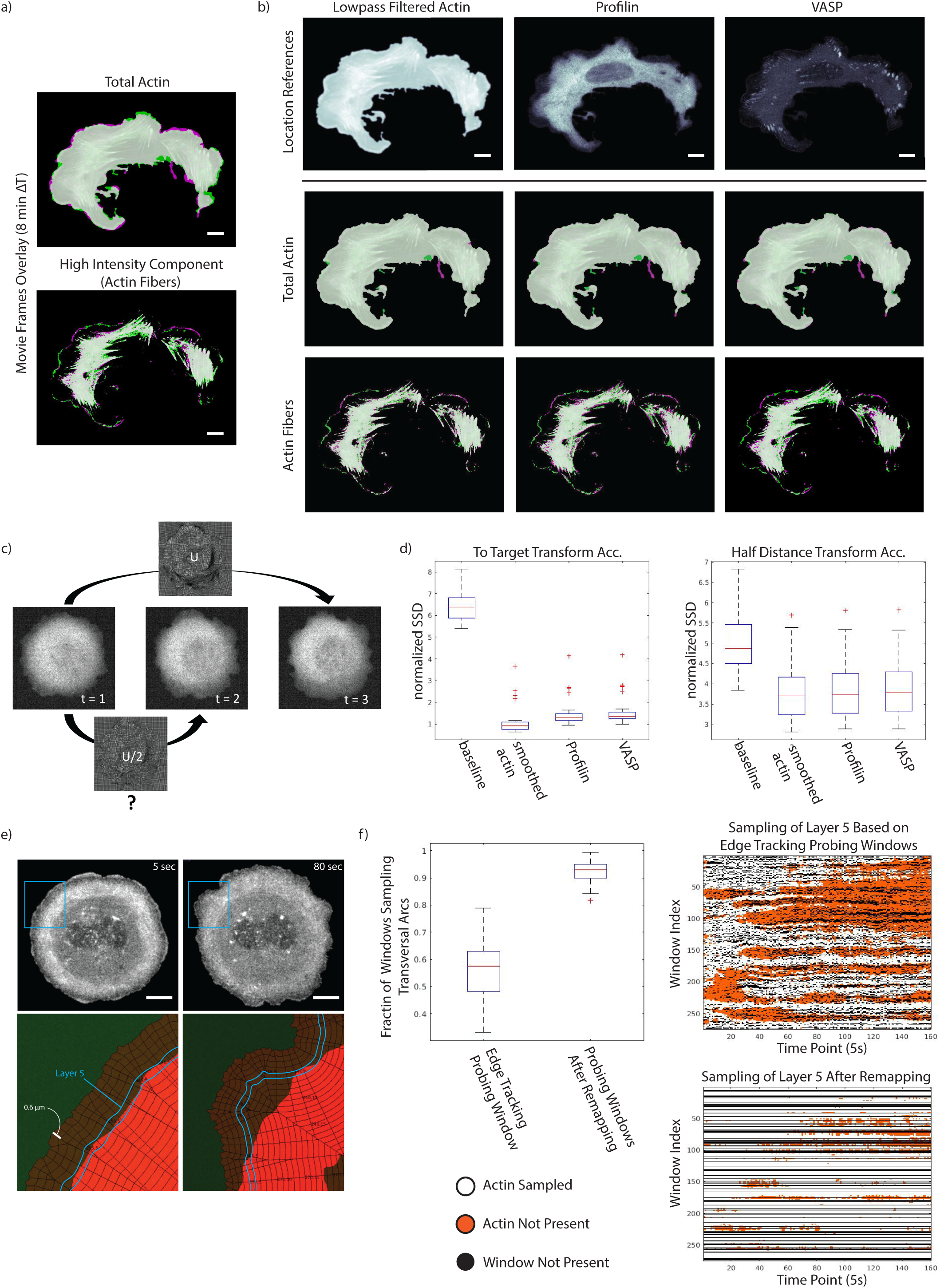
Impact of location fiducial on remapping accuracy. a) Overlay of two frames separated by 8 minutes of the total actin channel (top) and a high intensity component (defined by manually selected intensity cutoff) emphasizing actin fibers (bottom). b) Qualitative comparison of the remapping quality for both actin signals of interest using the lowpass filtered actin image (top row left), profilin (top row middle), and VASP as location fiducials. There were minimal differences in the resulting registration quality in both the Total actin (middle row) and actin Fiber (bottom row) components of the actin signal of interest. c) Diagram of the half-distance transform used as a ground truth for performance evaluation. First, we estimate the deformation on in diffeomorphic map from an input frame to a target frame, while skipping one frame in between. We then asked how well a half deformation map matches the middle frame under the expectation that intermediaries between original input and target follow a linear deformation path. d) Quantification of subcellular remapping accuracy using the half-distance transform ground truth. As expected, using the lowpass filtered actin signal as the location fiducial resulted in the most accurate match (Sum of squared distance SSD between target and remapped images) in both the to-target-transform (left) and half-distance-transform (right) ground truth scenarios. Overall, there were only small differences in accuracy between the choices of location fiducials. e, f) Comparison of full cell diffeomorphic mapping vs. edge tracking probing windows in sampling time courses of subcellular signals of interest. e) Snapshots of the actomyosin organization in a U2OS cell at an early (top left) and later (top right) stage before symmetry breaking. The cell displays characteristic transversal arcs behind a thin lamellipodia layer at the periphery. Towards the symmetry breaking event, the arcs begin to dissolve. In between, the transversal arcs follow the displacement of the cell edge. We detected these arcs using an intensity filter (red regions bottom row). Overlaid to the segmented arc region is a grid of 0.6 μm wide (at initial time point) and 0.6 μm deep probing windows, which follow the cell edge. Highlighted in blue, 5^th^ layer of probing windows, which sample the transverse arcs in early time points (left). In late time points the windows, which track with the cell edge movement fall outside the arc region. f) Sampling of the transverse arcs by the 5^th^ layer of probing windows tracking the cell edge or fixed after remapping (left). Space vs time heatmaps of the samples (right, top & bottom). The maps show the presence (white) or absence (orange) of transverse arcs. Windows tracking edge motion have frequent “drop-out” (black) in layers further away from the cell periphery. For direct comparison of indexed windows, we applied the reference time point probing windows to all frames in the remapped movie. “drop-out” windows (black) are persistent in time with remapping approach. All scale bars 10 μm.

Fig. 3 presents these metrics for four scenarios compiled from a movie of a U2OS cell simultaneously imaged for mNeonGreen (mNG)-actin, SNAP-profilin and Halo-VASP. To generate this test movie, we resampled the movie such that visible motion exists between any two given time points (1 frame/50 sec). Profilin is a cytoplasmic binding partner of monomeric actin and in concert with actin polymerases including formins and VASP, serves as a pacemaker for actin elongation [22–24]. VASP in addition is a component of the focal adhesion complex and presents a punctate cytoplasmic signal [25]. We used profilin, VASP, as well as down-sampled (Gaussian filtered with a sigma of 20 pixels) actin as location fiducials to remap the raw actin organization as the signal of interest. Like in Fig. 1, we chose again actin because of the dynamic and multi-factorially regulated structure, making it unlikely that any of the fiducial would fully capture the evolution over time. For a baseline we included the sum of squared intensity differences between unregistered original images over two frames (background set to 0). Both profilin and VASP as location fiducials greatly outperformed the baseline deviation (Fig. 3b), despite the diffuse image character in the former and the relatively scarce punctate pattern in the latter. Unsurprisingly the highest remapping accuracy was achieved with down sampled actin as the location fiducial. This shows that for generating precisely stabilized images of cytoskeleton structure in a reference frame our algorithm works best using a low pass filtered copy of the original signal for alignment.

Quite unexpected, the punctate VASP pattern produced nearly as good deformation maps for signal remapping as the continuously defined, diffuse profilin distribution. Upon close inspection, we can see only small differences in the remapping of subcellular actin structures between the first and last frames of a 20 min movie (Fig. 3a). We explain this with contributions made by the faint and diffuse but still spatially informative background pattern towards the estimation of the deformation field in between the salient puncta in the foreground. For the half-distance transform accuracy, we chose to skip only one frame since we were more likely to sample a steady state process during short time spans. Overall, the half-distance transform accuracy is worse than the to-target transform accuracy.(Fig. 3d) This is mostly attributable to poor registration near the cell edge, which moves on a much faster timescale than the cell interior [1, 21].

### Evaluation of Algorithm for the Extraction of Subcellular Time-Series

To further establish confidence that the remapping algorithm permits extraction of meaningful time-series throughout the cell, we compared the performance against our well-established cell peripheral windowing method [1, 8]. In brief, the windowing method tracks the cell edge and divides the cell into volumes indexed by radial position and depth in layers. While the proposed new registration-based approach can handle a large variety of cell morphodynamics, the windowing strategy has largely been used against stationary cell edge dynamics. We, therefore, chose a less motile U2OS cell (labeled with mNG-actin, Halo-CAAX, SNAP-profilin) undergoing stereotypical cell spreading to compare between the two methods.

In U2OS cells the periphery is demarcated by distinct and persistent network of transversal actin arcs. We segmented the arcs throughout the movie using a simple intensity threshold and hole-filling operation (Fig. 3e). We then applied the windowing algorithm to define layers of edge-tracking probing windows, each of which is 0.6 μm deep (Movie 4). In early time points the arcs begin ∼3 μm away from the cell edge (sampled by layers 5 to 8). We chose to examine the sampling of the circumferential actin region by layer 5 in a movie with edge-tracking windows and in a movie where the actin signal of interest was first remapped based on the CAAX membrane marker as location fiducial. In the remapped movie we applied the window positions from the reference frame to the entire movie as a stationary probing grid. Due to the fixed one-to-one window correspondence between layers, deeper layers exhibit drop-out events (Fig. 3f black streaks on the space-time representation of layer 5). Importantly, these drop-out windows do not persist over time since they are dependent on the geometry of the cell edge. In contrast, the window grid in a remapped movie has persistent drop-outs. The band between cell edge and transversal arcs, i.e. the lamellipodium, varies in width over time and also in space. Accordingly, in a sampling approach that preserves a constant distance from the edge, windows at the transition between lamellipodium and arcs alternate in the structure they sample. In contrast, the proposed remapping approach accounts for the variation in subcellular structures. Indeed, while edge-tracking windows in layer 5 sampled the transversal arc structure in only 58% of the windows and time points, 92% of the stationary windows of layer 5 in the remapped movie sampled transversal arcs (Fig. 3f barplot). The loss of connection to subcellular structures has been a serious limitation for many studies relying on edge tracking windows as they sample time series associated with distinct regulatory regimes [26, 27]. The proposed remapping resolves this issue now for any subcellular structure that follows the diffeomorphism defined by the location fiducial.

### Time-Series of Profilin reveal Spatiotemporal Organization of Dynamic Concentrations

Using windows sampled in a narrow band along the cell edge, previous work in our lab has shown that molecular signaling activities implicated in cytoskeleton regulation can be spatially partitioned into micro-domains of similar dynamics [8]. Equipped with a tool for sampling time series across an entire cell, we hypothesized that the same principle of spatial coherence in dynamic behavior could be applied to map out the subcellular organization of protein dynamics. Exploring this possibility, we examined the subcellular organization of profilin.

Profilin is a small molecule binding partner of monomeric actin and profilin-actin is considered the physiological substrate of filament growth [23, 24, 28]. Numerous biochemical experiments have shown specific interactions between profilin-actin and key cytoskeletal regulators (such as formins) suggesting that distinct actin structures would colocalize with local pools of profilin with distinct dynamics [22, 29]. However, endogenously SNAP-tagged profilin in live cells displayed a diffuse signal with no visually discernable pattern beyond intensity variations that related to the integration along the optical axis of fluorescence in variably thick cells (Fig. 4a-c, left). To discover patterns of dynamics, we measured in each pixel of the remapped cellular footprint the statistical coherence of the time-series within a 3×3 neighborhood (Fig. 4a-c, right). Coherence has a value between 0 and 1, where higher values indicate self-similarity among the 9 time-series. This analysis resembles fluorescence correlation spectroscopy [30]. However, at the time scales of our movies, the coherence is likely not due to a physical property but due to the cell actively maintaining such local similarity through either biochemical interactions or locally constrained morphodynamics.

**Figure 4.**
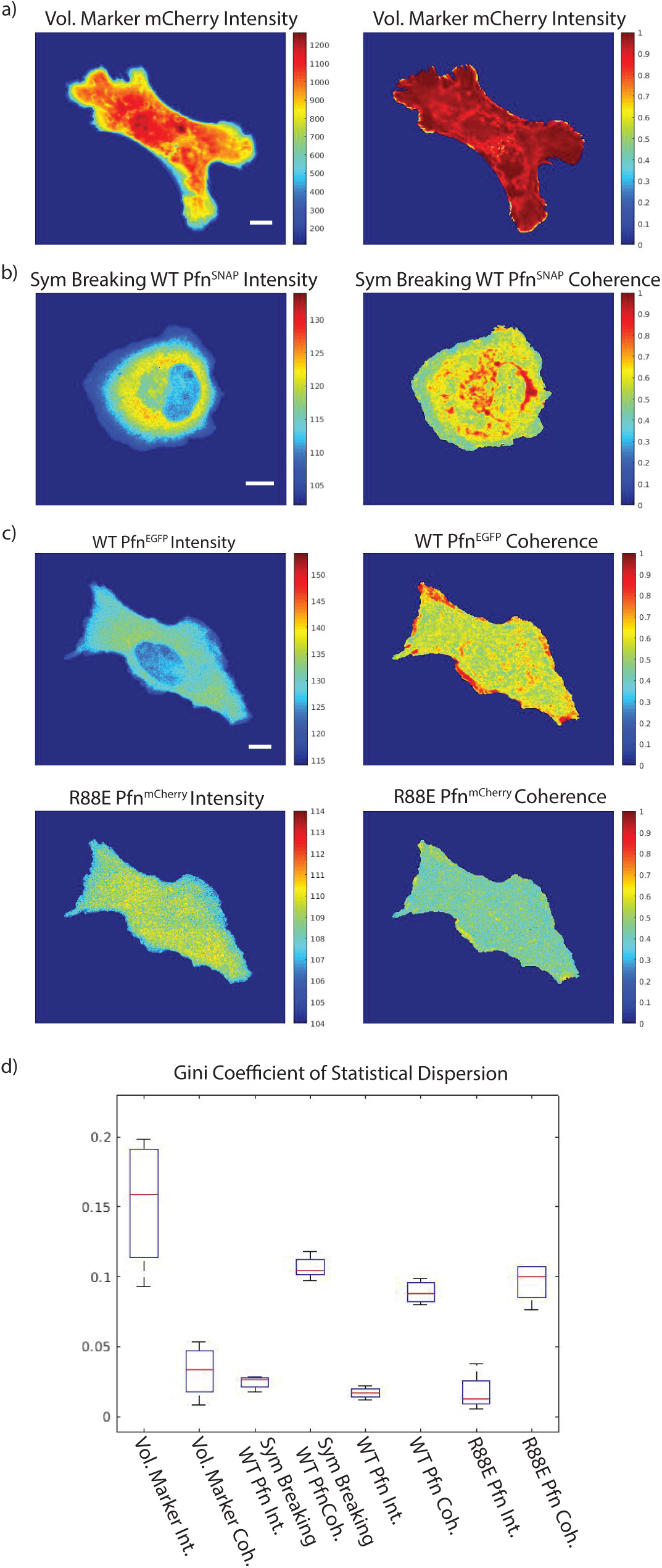
Profilin dynamics in live cells show patterns in local time series coherence. Raw intensity of diverse profilin probes vs coherence of the profilin probes’ time series in a U2OS cell. The computation of reliable coherence scores is enabled by the remapping of the movie into a stable reference frame. a) Control experiment using an mCherry cytoplasmic volume marker. As expected, coherence of a diffuse signal is high across the entire cell footprint. For remapping, a Halo-CAAX tag membrane marker was used as the location fiducial. b) Symmetry breaking cell expressing SNAP-profilin at endogenous levels. For remapping, using a Halo-CAAX tag membrane marker was used as the location fiducial. c) WT EGFP-profilin (top) and R88E mApple-profilin mutant (bottom) concurrently expressed at low concentration from a leaky CMV100 promotor in profilin knockout cell. For remapping, lowpass filtered actin signal was used as the location fiducial. WT and R88E profilin display distinct coherence patterns. d) Quantification of subcellular heterogeneity of the signals observed in a-c using the Gini coefficient. All scale bars 10 μm.

We performed this analysis on 3 cell populations: First, U2OS cells expressing cytoplasmic mCherry as a volume marker in a background of mNG-actin and Halo-CAAX (Fig. 4a). Second, U2OS cells expressing fluorescent endogenously SNAP-tagged profilin, mNG-actin, and Halo-CAAX (Fig. 4b). These cells were further treated with the myosin-II inhibitor Blebbistatin to induce symmetry breaking throughout the movie, as described by Lomakin et al [31]. These cells allowed us to follow the changes in profilin dynamics in response to a change in global cell morphology. Third, we exogenously co-expressed wild-type s EGFP-profilin (WT) and a mutant mApple-profilin (R88E) at very low levels using a leaky CMV100 alongside SNAP-actin in profilin KO cells (Fig. 4c). The R88E mutation abrogates binding to actin [29] therefore we hypothesized that the dynamics of the mutant profilin would follow a different coherence pattern than that of the wildtype. In the first 2 cases, we remapped the movies on a central reference frame using the CAAX marker as a location fiducial (images not shown). Due to the limitations in expressing yet another tag for live cell imaging, we remapped the double profilin labeled cells using a down-sampled actin channel set to mimic the results of CAAX based remapping.

The coherence analysis revealed drastically different organizational patterns relative to the raw profilin signal. High coherence is observed in select sites around the cell edge, in puncta throughout the cytoplasm, and around the nucleus in both the endogenously and exogenously tagged profilin. We quantified the increase in heterogeneity produced by application of the coherence operator via the Gini coefficient of statistical dispersion (Fig. 4d). The Gini coefficient occupies a value range between 0 to 1, where 1 indicates a signal concentrated in one pixel of the cell and 0 indicates homogeneous signal across all pixels of the cell [32]. For all three cell conditions, the coherence of the profilin intensity is more heterogenous than the raw profilin signal, indicating a high level of local dynamic organization of profilin, potentially related to its roles in facilitating actin polymerization. As a control, we performed the same analysis for a cell expressing mCherry as a volume marker. In this case the intensity shows a higher level of heterogeneity than the coherence value, because of significant variation in cell thickness. The coherence of the mCherry volume marker was homogeneously high throughout the cell.

### Profilin Coherence is Related to Actin Dynamics

Since WT and R88E profilin displayed distinct coherence patterns in the same cell, we hypothesized that WT Profilin coherence would show a stronger relationship with actin dynamics. Our coherence calculation thus far relied on time-series spanning the entire movie. To test whether profilin coherence changes in concert with actin dynamics we computed a coherence time-series using a moving window of 25 frames, i.e. 1/10 of the length of our shortest movies (Fig. 5c, Fig. 5d, Movie 5, and Movie 6). The resulting time series could then be locally correlated with a measure of the actin signal fluctuations. To match the time scales of changes in profilin coherence and actin dynamics, we computed the entropy of the actin signal at every location of the cell using the same moving window approach (Movie 7). This signal transform extracts from the overall fairly static actin images (Fig. 5a) second-to-minute scale processes such as stress fiber growth and movements and retrograde flow (Fig. 5b).

**Figure 5.**
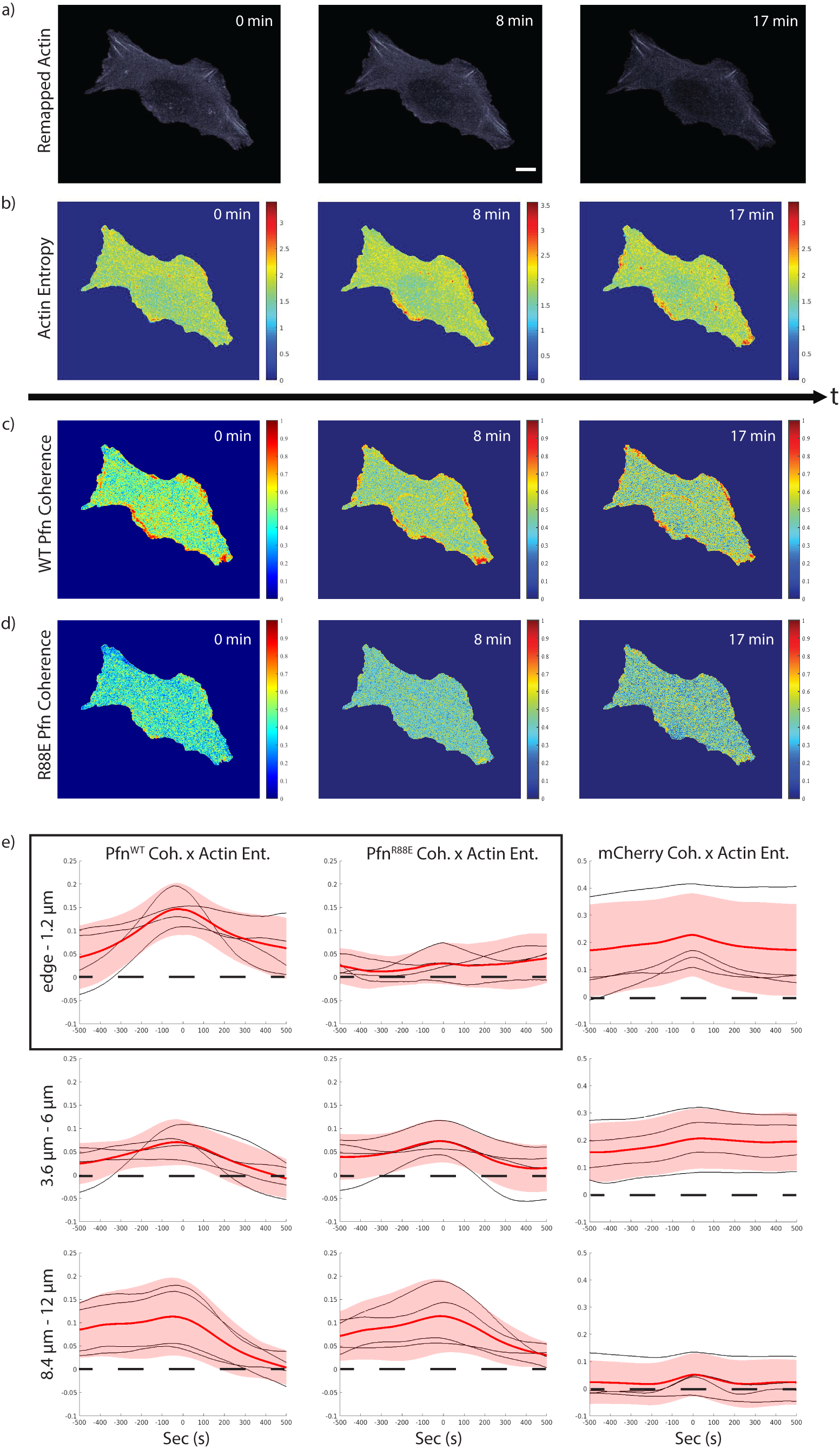
Co-expressed wildtype and mutant Profilin exhibit different relationship to Actin dynamics. a-d) Signal transforms to relate profilin organization to actin dynamics. The transforms are enabled by the remapping of a movie to a stable reference frame. The data displays U2OS cells expressing actin-SNAP (a), with the entropy of the actin signal shown in (b), WT EGFP-profilin (c), and R88E mApple-profilin (d). The remapping was accomplished using lowpass filtered actin as a location fiducial. To reduce perturbation by overexpression, the two profilin constructs were expressed in a Profilin null background. Panels in (c) and (d) display the dynamics of the diffuse image signals as the spatially local coherence over a rolling window of 25 time points. The SNAP-actin signal consists of various filament forms as well as a diffuse background of monomers. The mixture makes a direct cross correlation to other molecular processes difficult to interpret. We therefore extracted the entropy over a rolling window of 25 time points (250 s), which indicates relative stability of actin structures (high entropy delineates regions of high polymer turnover or translocation). e) Cross correlation of actin entropy and profilin coherence in subcellular regions defined by distance from the cell edge. We performed this analysis on 3 zones: 1.2 μm from the cell edge (top row) roughly corresponding to lamellipodia, 3.6 – 6 μm from the cell edge (mid row) roughly corresponding to transverse arcs, and 8.4 – 12 μm corresponding to regions around the cell nucleus. As a control we also calculated the cross correlation between actin entropy and the coherence of an mCherry volume marker (right column). WT profilin and actin show a significant correlation only in the band 0 – 3.6 μm from the edge, confirming the biochemical function of profilin as a promoter of actin polymerization. The correlation collapses for R88E profilin, which is deficient in actin binding (boxed). The coupling between profilin and actin dynamics is absent in regions more distal from the cell edge. All scale bars 10 μm.

We then examined the cross correlation of profilin coherence and actin entropy near the cell edge (edge – 1.2 μm), around the circumferential actin network (3.6 – 6 μm), and around the nucleus (8.4 – 12 μm) since the data in Fig. 4 indicated patterns of high profilin coherence in these positions (Fig. 4c). As expected, near the cell edge we observed higher cross correlation between actin entropy and WT profilin when compared with R88E profilin (Fig. 5e boxed). The difference is less substantial in deeper regions of the cell because both actin entropy and profilin coherence are temporally less salient. This is reinforced by the fact that the volume marker’s cross correlation with actin entropy is high in the periphery but low near the nucleus where there is a persistently high volume marker coherence but rare high actin entropy events (Fig. 5e). Together, these analyses indicate how the proposed remapping of signals to a static reference cell geometry permits the application of time series preprocessing in order to extract subtle but significant dynamic parameters in otherwise diffuse signals that for the purpose of assessing the local interactions between molecular processes.

### Profilin Coherence Correlation with Actin Dynamics is Responsive to Perturbation

To test the hypothesis that the observed relationship between profilin coherence and actin entropy relates to profilin’s functions as a modulator of actin polymerization, we analyzed cells undergoing symmetry breaking (Movie 8). Symmetry breaking is a process where the cell transitions from a stable rounded state to a polarized migratory state. In doing so the cell must greatly reorganize its actin network. After remapping the entire movie to a common reference frame just before the symmetry breaking event (Fig. 6a, see also Fig. 1b), we split the movies into before and after symmetry breaking section to compute actin entropy (Fig. 6b) and profilin coherence (Fig. 6c). Comparing the spatial distribution of actin entropy before and after induction of symmetry breaking by Blebbistatin, actin entropy initially increases throughout the entire protrusive front (Fig. 6b panels 0 min vs 4 min) as the cell begins to repolarize before decaying again to remain elevated just along the cell edge (Fig. 6b panels 17 min and 21 min). This is consistent with the notion that the high actin turnover, corresponding to membrane ruffles and transversal arcs all around the cell edge, is shifted during symmetry breaking towards the wide lamellipodia and lamella regions at the new cell front [31]. This reorganization of actin dynamics is paralleled by a reorganization of high profilin coherence (Fig. 6c). Specifically, before symmetry breaking, we see high profilin coherence right along the cell edge, on top of transversal arcs and around the nucleus. We see background levels of coherence near the rear retraction fibers. There exists a faint but noticeable low coherence band between the leading edge and transversal arcs. Again, locally high profilin coherence indicates high spatial coupling in the concentration fluctuations of profilin-actin complexes that are fed into actin polymers by nucleators such as formins which promote the growth of linear actin structures [23, 24, 28]. Importantly we do not expect these zones to be an artifact of the remapping process as their size significantly exceeds the approximate deformation of the movie frames. The facts that R88E and WT profilin exhibit clearly different dynamics at the cell edge (Fig. 5c) and that the more stable cell rear where we expect lower actin turnover does not exhibit higher coherence leads us to conclude that our data report effects driven by the profilin-actin interaction . Hence, this analysis – enabled by the proposed remapping algorithm – visualizes for the first time directly in a living cell profilin’s function and organization as a critical facilitator of actin assembly.

**Figure 6.**
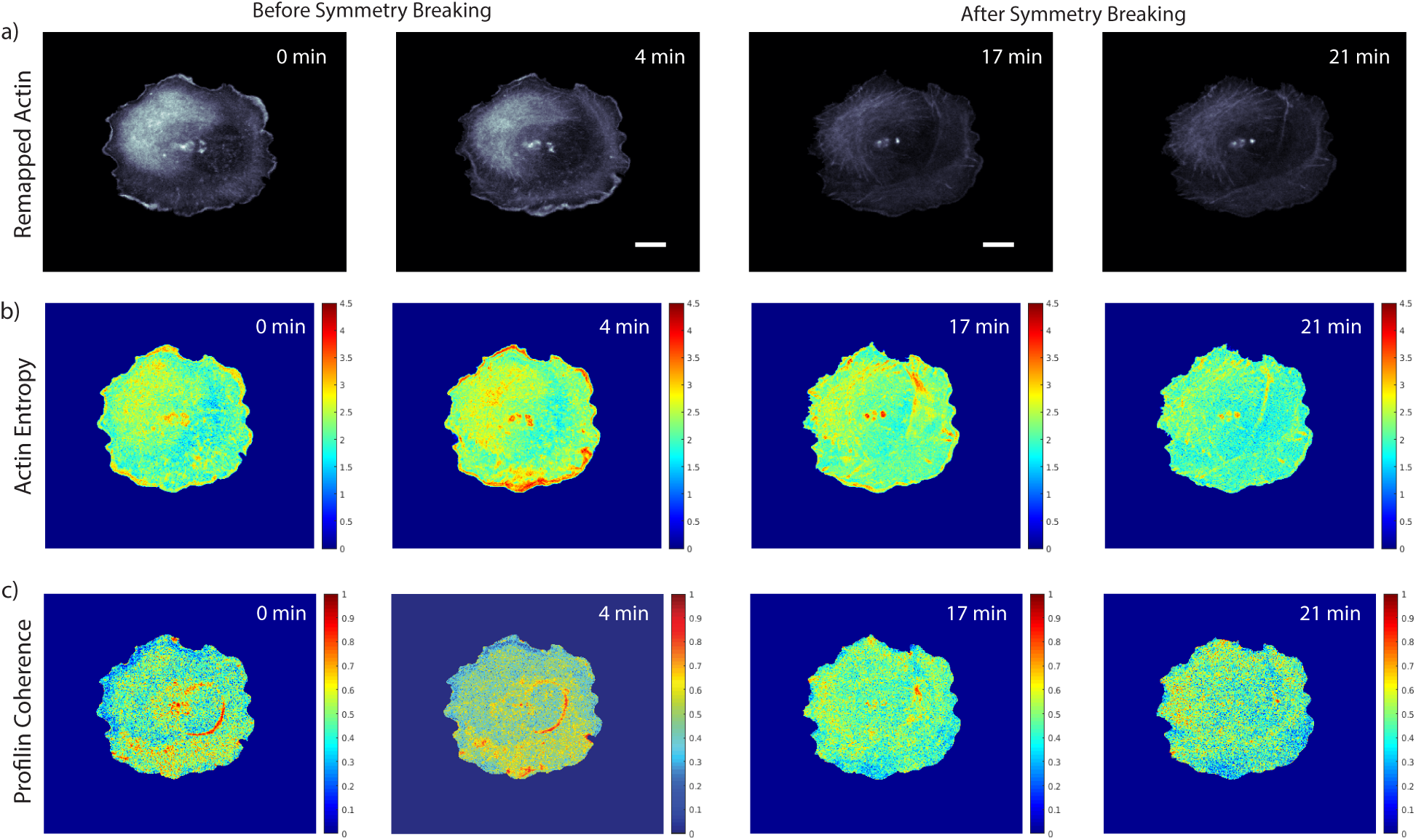
Drug induced symmetry breaking reveals polarization dependent organization of profilin coherence. a) mNG-actin signals before and after symmetry breaking in U2OS cells induced by 25 μM Blebbistatin. The remapping of the movie to a reference frame was accomplished using Halo-CAAX as a location fiducial. Symmetry breaking occurs at approx. 10 min after imaging. We designated before symmetry breaking as the time span from start to 4 min and after symmetry breaking as 17 min to end of imaging. b) Actin entropy over a rolling window of 25 time points before and after symmetry breaking. c) Profilin coherence over a rolling window of 25 time points before and after symmetry breaking. Both the actin entropy and profilin coherence reveal a shift in high values away from the cell front to the cell center and back during the symmetry breaking process showing that profilin organization is responsive to changes in subcellular actin polymerization. All scale bars 10 μm.

## Discussion

In this work we implemented a non-linear image registration framework to analyze subcellular protein dynamics in cells undergoing substantial morphological variation throughout the observation window. The key contribution of our work to the sizable literature on non-linear image registration algorithms is the capacity to handle the high noise and small structures of interest present in cell microscopy. This was accomplished by introducing a cellular motion mask as a regularization term in the image mapping objective function and by enforcing diffeomorphism via a sorting of the displacement field. The former causes the map estimator to bypass local minima, the latter permits the preservation of fine grained structures during the mapping.

We demonstrated the capacity of this framework by extracting biological insight through an observational study of previously uninterpretable signals. Importantly, the algorithm supports the extraction of reliable time-series at every pixel position in the cell footprint and thus enables studies of previously inaccessible spatiotemporal organization of molecular processes deep in a cell. Such analyses were hitherto limited to a narrow rim along the cell boundary, where time series could be sampled by a deformable window grid following the edge motion. We illustrated these new features by examining the relationship between profilin and actin dynamics. Profilin lends a highly diffuse image signal with visually uninterpretable variation. Actin lends a mixture of highly structured and amorphous image signal components. As a cell undergoes morphological changes, the structured components often display complex patterns of deformation whereas the amorphous components undergo often difficult to follow flows. Because of our proposed image registrations, we were able to transform both the profilin and actin signals into secondary signals that readily revealed the spatiotemporal coupling of profilin and actin dynamics, even during a cellular symmetry breaking event that produces large scale cell shape changes.

While designing this framework we expected the resulting time series to strongly depend on the choice of the location fiducial used for the mapping estimation. We were surprised that a punctate signal (VASP) permits the algorithm to remap actin structures over time with an accuracy that is comparable to the mapping accuracy supported by a location fiducial derived from a diffuse signal like profilin or a blurred version of the actin signal itself. This is likely because adhesion proteins like VASP have a faint but implicitly traceable diffuse component that constrains the map estimator alike profilin and actin signals. Moreover, cytoskeleton structures are highly coordinated in healthy cells and the cytoplasm is a dense compartment meaning that subcellular molecular flows in general are coupled. Hence, there is some degree of tolerance in choosing a location fiducial.

Most importantly our framework opens the door to analyses of subcellular signals with dynamics that occur on the same or slower timescale as cell morphological changes. For processes much faster than cell morphological changes (e.g. cell electrical potentials and calcium signaling), microscopy has produced predictive quantitative models for *in vivo* signaling behavior and outcomes because cellular changes could be ignored. However, the majority of subcellular activities likely occur on a timescale that matches cell morphodynamics. While existing approaches have generated anecdotal and often visually-guided analyses of these processes in select cell regions, our framework now supports an unbiased analysis across the entire cell. This is the starting point for robust pattern recognition in cell biological activity.

## Materials and Methods

### Plasmids

pSpCas9(BB)-2A-GFP (PX458) and pX335-U6-Chimeric_BB-CBh-hSpCas9n(D10A) were gifts from Dr. Feng Zhang (Addgene plasmid #48138 and #42335, respectively). Gene-targeting single guide RNAs (sgRNAs) were designed using the online program CRISPor (http://crispor.tefor.net) [33]. pmCherry-C1 was from Clontech. The self-cleaving vector pMA-tial1 was from Dr. Tilmann Bürckstümmer [34]. pBlueScript II SK(+) was from Agilent. The primers to clone the profilin-1 (PFN1)-targeting sgRNA pair used for knockouts were 5’- CACCGTCGATGTAGGCGTTCCACC-3’ and 5’-AAACGGTGGAACGCCTACATCGAC-3’ and were cloned into PX458. PFN1-targeting sgRNA pairs for the knock-in were 5’- CACCGGCTGCTACTGGGGCTGCTCTCGG -3’, 5’- AAACCCGAGAGCAGCCCCAGTAGCAGCC -3’, 5’- CACCGCGCCTACATCGACAACCTCATGG -3’ and 5’- AAACCCATGAGGTTGTCGATGTAGGCGC -3’, all cloned into PX335. The self-cleaving donor vector containing the Blasticidine selection cassette was previously described [35, 36]. The N-terminal SNAP-tag knock-in donor was flanked with homologous arms targeting the first exon of PFN1 (BAC library: CH17-25E2) including a 13 amino acid linker (SGRTQISSSSFES) in between SNAP-tag and PFN1, a configuration well-characterized to preserve all functional properties of profilin-1 [37, 38], and cloned into pBlueScript II SK(+). pLVXCMV100- mNeonGreen-18-actin and pLVXCMV100-Halo-21-VASP were previously described[26]. pLVXCMV100- Halo-CAAX was generated by Gibson Assembly (NEB) using HaloTag sequence as a template and the following primers: 5’-attaactagtgccaccatggcagaaatcggtactggctttcc-3’, 5’- ttacataattacacactttgtctttgacttctttttcttctttttaccatctttgctcatcttttctttatgGCCGGAAATCTCGAGCGTCGAC-3’ and 5’-attacgcgtTTACATAATTACACACTTTGTCTTTGACTTCTTTTTCTTC-3’.

N-terminally tagged EGFP-C-profilin-10 and mApple-C-profilin-10, both harboring mouse profilin-1 and a 10 amino acid linker (SGLRSRAQAS) were gifts from Michael Davidson (Addgene plasmids #56438 and #54940). The actin-binding deficient profilin-1 mutant R88E [39] was generated by mutating the Arginine at position 88 on profilin to Glutamic acid (R88E) using primers 5’-gagACCAAGAGCACCGGAGGAGCCCC-3’ and 5’-AAGATCCATTGTAAATTCCCCGTCTTGCAGCAGTG-3’ and PfuUltra II Fusion High-fidelity DNA polymerase (Agilent). EGFP-profilin and mApple-profilin were subsequently subcloned into the SpeI/MluI sites of the lentiviral vector with attenuated CMV promoter, pLVXCMV100 [40], using primers 5’- ttaactagtGCCACCATGGTGAGCAAGGGCGAG -3’, 5’- ttaactagtgccaccATGGTGAGCAAGGGCGAGGAGAATAACATGG -3’ and 5’- aaacgcgtTCAGTACTGGGAACGCCGCAGGTGAGA -3’. All newly generated constructs were sequence verified.

### Antibodies

Mouse monoclonal anti-profilin-1 (Santa Cruz; B-10; sc-137235), mouse monoclonal anti-vinculin (Sigma; V9264) and mouse monoclonal anti-Actin (Sigma; AC-15; A1978) antibodies.

### Cell Lines

Human Osteosarcoma U2OS cells were cultured in DMEM media supplemented with 10% fetal bovine serum (Sigma; F0926-500ML) in a humified incubator at 37 °C and 5% CO2. All cells were tested for mycoplasma using a PCR-based Genlantis Mycoscope Detection Kit (MY01100).

Lentiviral particles were generated using the packaging vectors psPAX2 and pMD2.G (Addgene plasmids #12260 and #12259). Infected cells were bulk sorted using FACS.

Profilin-1 (PFN1) knockout U2OS cells were generated by co-transfecting the PFN1-targeting sgRNA supplemented with the self-cleaving Blasticidine selection cassette. Genome-edited cells were selected using 5 µg/ml Blasticidine S selection (Thermo) and isolated using 8 mm colony cylinders (Sigma). Knockouts were verified using western blot with mouse anti-Profilin-1 antibodies. Profilin KO clone #2 were used for subsequent experiments (Supplemental Fig 1).

Endogenously SNAP-tagged profilin-1 cells were generated by co-transfecting the two pairs of PFN1-targeting sgRNAs with Cas9 nickase supplemented with the donor vector. Genome-edited cells were labeled with SNAP-Cell Oregon Green (S9104; NEB) and single cell sorted into 96 wells plated coated with attachment factor (S006100; Gibco). Successful genome-edited cells were validated using western blotting.

mCherry was expressed in U2OS cells expressing mNeonGreen-actin and Halo-CAAX by transient transfection with polyethylenimine and 1 µg of pmCherry-C1 one day prior to imaging. U2OS profilin-1 KO U2OS cells were rescued with wild-type EGFP-profilin and actin-binding deficient mApple-profilin mutant (R88E) using a lentiviral construct driven by a truncated CMV promoter as discussed above. Infected cells were bulk sorted using FACS.

### Live Cell Imaging

Cells were counted using Cellometer Auto 1000 Bright Field Cell Counter (Nexcelom) and 100.000 cells were seeded on 10 µg/ml fibronectin-coated glass bottom 35 mm. Halo-CAAX was labeled using Janelia Fluor 549 HaloTag ligand (200 - 400 nM; GA1110; Promega) and endogenous SNAP-Profilin-1 were labeled using SNAP-Cell 647-SiR (500 nM – 1 µM; S9102S; NEB), 30 minutes prior to imaging. Cells were imaged in phenol-red free DMEM supplemented with 20 mM HEPES pH 7.4 on a climate-controlled (maintained at 37 oC), fully motorized Nikon Ti-Eclipse inverted microscope equipped with Perfect Focus System, an Andor Diskovery illuminator coupled to a Yokogawa CSU-XI confocal spinning disk head with 100 nm pinholes, and a 60x (1.49 NA) APO TIRF objective (Nikon) with an additional 1.8x tube lens, yielding a final magnification of 108x (Andor Technology). Images were recorded at 3 Hz frame rate using a scientific CMOS camera with 6.5-µm pixel size (pco.edge).

### Blebbistatin Treatment to Induce Symmetry Breaking

Cells were treated with 25 μM myosin II inhibitor blebbistatin in DMEM for 5 min prior to imaging as per [31]. Drug treatment was offset to allow for a complete 20 min imaging run per dish under the expectation that symmetry breaking would occur at approx. the 10 min mark. Fields were selected manually to center a single individual cell and cells were selected for a flat rounded morphology. After imaging, the movies were examined for spontaneous symmetry breaking near the midpoint. We utilized the first 1/3 of the movie as before symmetry breaking samples and the last 1/3 of the movies as post symmetry breaking samples.

### Remapping Pipeline Parameters

For all published results we set **α** = 1 for a maximum step size of 1 pixel. For sequentially transformed movies for time series analysis we set **n** = 100 iterations per time step and **Kdiff** = 1.5 pixels. For transforms between the first and last image frames we set **n** = 2000. For ½ deformation accuracy we set **n** = 200.

### Profilin Coherence Analysis

For every subcellular location we sampled a 3×3 matrix of 9 time series centered on the target pixel. We calculated the correlation coefficient for all 72 =non-identical time-series pairs, where the correlation coefficient ***ρ*(A, B)** between time series **A** and **B** is:

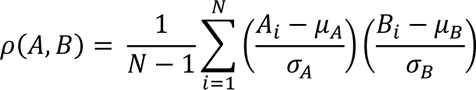

**μ_A_** and **σ_A_** are the mean and standard deviation of **A**, respectively, **μ_B_** and **σ_B_** are the mean and standard deviation of **B**, and **N** is the number of time points in the series **A** and **B**. The coherence is the mean of resulting coefficients. For edge positions we removed the out of the cell time series from the initial 3×3 matrix. We calculated coherence over the course of the entire movie and in 20 frame moving windows (1/10 length of typical movie) to calculate coherence change over time.

### Actin Entropy Calculation

The Actin channel contains a mixture of local monomers, branched actin, and linear bundles making correlational analysis prohibitively difficult. To address this, we calculated the Shannon information entropy for subcellular actin intensity in 20 frame moving windows (1/10 the length of a typical movie). For every actin intensity series **X** the entropy of said series **H(X)** is:

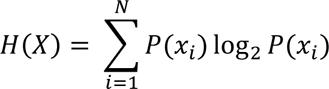

Where **x_i_** is a value in series **X** and **P(x_i_)** is the probability of drawing **x_i_** from series **X**. In effect this reveals the instability of local actin and captures both concentrations driven, and polymer type driven intensity shifts in a single value.

### Windowing Parameterization

In the application of the published algorithm [1, 8], we used the option ‘Constant number’ as a method of propagating the windows to the next time frame. To select the size of the probing windows, we reviewed the past applications of the approach to determine a representative setting. This determined the size of the probing windows to be 600×600 nm^2^ in parallel and perpendicular to the edge. Over time and depth, the lengths parallel to the cell edge are allowed to vary but the lengths perpendicular to the edge are fixed.

## Acknowledgement

We thank Dr. Dick McIntosh (University of Colorado, Boulder, CO) for providing the U2OS osteosarcoma cells, Dr. Tilmann Bürckstümmer and all Addgene depositors for sharing reagents and Dr. Dana Reed (UT Southwestern Medical Center) for her logistical support and laboratory management. We also thank the UT Southwestern BioHPC facility for providing high-performance computing systems. This study was supported by R35GM136428 to GD.

## SUPPLEMENTAL MATERIALS

**Supplemental Figure 1.**
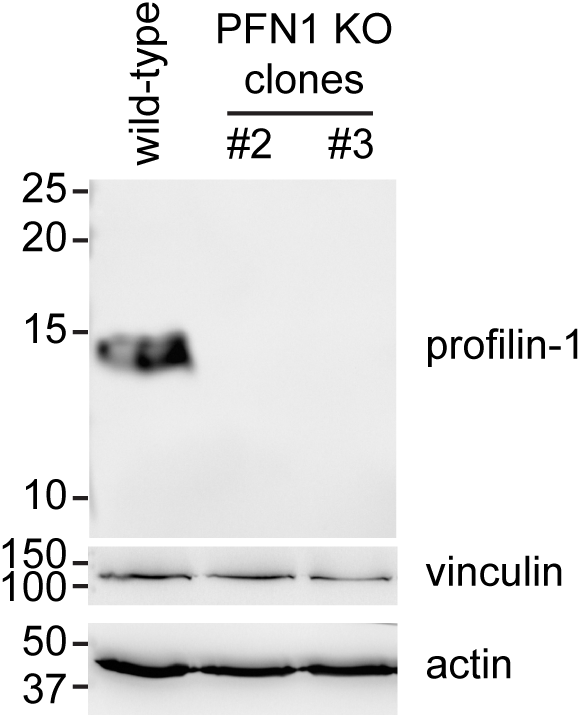
Verification of profilin knockout clones. Profilin knockout was verified using western blotting using mouse monoclonal anti-profilin-1 antibodies. Vinculin and actin provided as loading control. See materials and methods for antibody source.

**Movie 1. Comparison of original and remapped CAAX location fiducial movies**

Cell: U2OS SNAP-CRISPR-Profilin, mNG-Actin, Halo-CAAX treated with 25 μM Blebbistatin Left: Original movie

Right: Remapped movie Acquisition rate: 1 frame/5s Replay rate: 21 frames/s

**Movie 2. Comparison of original and remapped Actin movies**

Cell: U2OS SNAP-CRISPR-Profilin, mNG-Actin, Halo-CAAX treated with 25 μM Blebbistatin Top left: CAAX original movie

Top right: CAAX remapped movie Bottom left: Actin original movie Bottom right: Actin remapped movie Acquisition rate: 1 frame/5s

Replay rate: 21 frames/s

**Movie 3. Edge highlighted comparison of original and remapped Actin movies**

Cell: U2OS SNAP-CRISPR-Profilin, mNG-Actin, Halo-CAAX treated with 25 μM Blebbistatin Left: original movie

Right: remapped movie

Color scale: red-current time point edge position -> blue-past time point edge position Acquisition rate: 1 frame/5s

Replay rate: 21 frames/s

**Movie 4. Windowing of transverse arcs in symmetry breaking cell**

Cell: U2OS SNAP-CRISPR-Profilin, mNG-Actin, Halo-CAAX treated with 25 μM Blebbistatin Left: full cell view

Right: zoomed in view

Color: red-subcellular windows, pink-transverse arc detection, black lines-window boundaries Acquisition rate: 1 frame/5s

Replay rate: 7 frames/s

**Movie 5. Comparison of Profilin signal and Profilin Coherence**

Cell: U2OS SNAP-CRISPR-Profilin, mNG-Actin, Halo-CAAX treated with 25 μM Blebbistatin Left: Profilin signal

Right: Profilin coherence in 25 time point moving windows Acquisition rate: 1 frame/10s

Replay rate: 7 frames/10s

**Movie 6. Comparison of WT-Profilin and R88E-Profilin coherence in same cell**

Cell: U2OS EGFP-Profilin^WT^, mCherry-Profilin^R88E^, SNAP-Actin Left: WT-Profilin coherence in 25 time point moving

Right: R88E-Profilin coherence in 25 time point moving windows Acquisition rate: 1 frame/10s

Replay rate: 7 frames/10s

**Movie 7. Comparison of Actin signal and Actin entropy**

Cell: U2OS SNAP-CRISPR-Profilin, mNG-Actin, Halo-CAAX treated with 25 μM Blebbistatin Left: Actin signal

Right: Actin entropy Acquisition rate: 1 frame/5s Replay rate: 7 frames/5s

**Movie 8. Profilin coherence before and after symmetry breaking**

Cell: U2OS SNAP-CRISPR-Profilin, mNG-Actin, Halo-CAAX treated with 25 μM Blebbistatin Left column: before symmetry breaking

Right column: after symmetry breaking Top row: original Actin signal

Mid row: remapped Actin signal

Bottom row: Profilin coherence in 25 time point moving windows Acquisition rate: 1 frame/5s

Replay rate: 7 frames/5s

